# Two sets of wing homologs in the crustacean, *Parhyale hawaiensis*

**DOI:** 10.1101/236281

**Authors:** Courtney M. Clark-Hachtel, Yoshinori Tomoyasu

## Abstract

The origin of insect wings is a biological mystery that has fascinated scientists for centuries. Through extensive investigations performed across various fields, two possible wing origin tissues have been identified; a lateral outgrowth of the dorsal body wall (tergum) and ancestral proximal leg structures^1,2^. With each idea offering both strengths and weaknesses, these two schools of thought have been in an intellectual battle for decades without reaching a consensus^3^. Identification of tissues homologous to insect wings from linages outside of Insecta will provide pivotal information to resolve this conundrum. Here, through expression analyses and CRISPR/Cas9-based genome-editing in the crustacean, *Parhyale hawaiensis*, we show that a wing-like gene regulatory network (GRN) operates both in the crustacean terga and in the proximal leg segments, suggesting that (i) the evolution of a wing-like GRN precedes the emergence of insect wings, and (ii) that both of these tissues are equally likely to be crustacean wing homologs. Interestingly, the presence of two sets of wing homologs parallels previous findings in some wingless segments of insects, where wing serial homologs are maintained as two separate tissues^4–7^. This similarity provides crucial support for the idea that the wingless segments of insects indeed reflect an ancestral state for the tissues that gave rise to the insect wing, while the true insect wing represents a derived state that depends upon the contribution of two distinct tissues. These outcomes point toward a dual origin of insect wings, and thus provide a crucial opportunity to unify the two historically competing hypotheses on the origin of this evolutionarily monumental structure.

---

The identification of serially homologous structures can be a powerful approach to reveal the life history of complex structures, as serially homologous structures can undergo varying degrees of evolutionary change in different body parts (such as in different segments of insects)^8,9^. Recent molecular attempts to identify wing serial homologs in some wingless segments of insects have shed light on the insect wing origin debate^4–7,10^, some of which have suggested that wings have a dual origin and are formed from a combination of the two previously proposed origin tissues^4–7^. However, this approach is inherently limited to the lineages where wings have already evolved, preventing us from obtaining a comprehensive evolutionary history of this structure. Identifying homologous tissues between different taxa can circumvent this limitation by helping us reconstruct the tissues that were likely present in the common ancestor of these groups, thus providing crucial information on how novel structures arise. As a member of a sister group to insects, the crustacean, *Parhyale hawaiensis*, provides an excellent opportunity to broaden the search for wing homologs. *Parhyale* are a well-established crustacean model for evo-devo studies^11–13^ and their dorso-ventral (DV) body plan remains largely similar to that of insects. Both of the tissues that correspond to the two proposed wing origins, the dorsal terga and the proximal leg segments (Fig. 1a), are present in *Parhyale*, allowing us to evaluate the evolutionary relationship of these tissues to the insect wing and other wing serial homologs.

**Figure 1.**
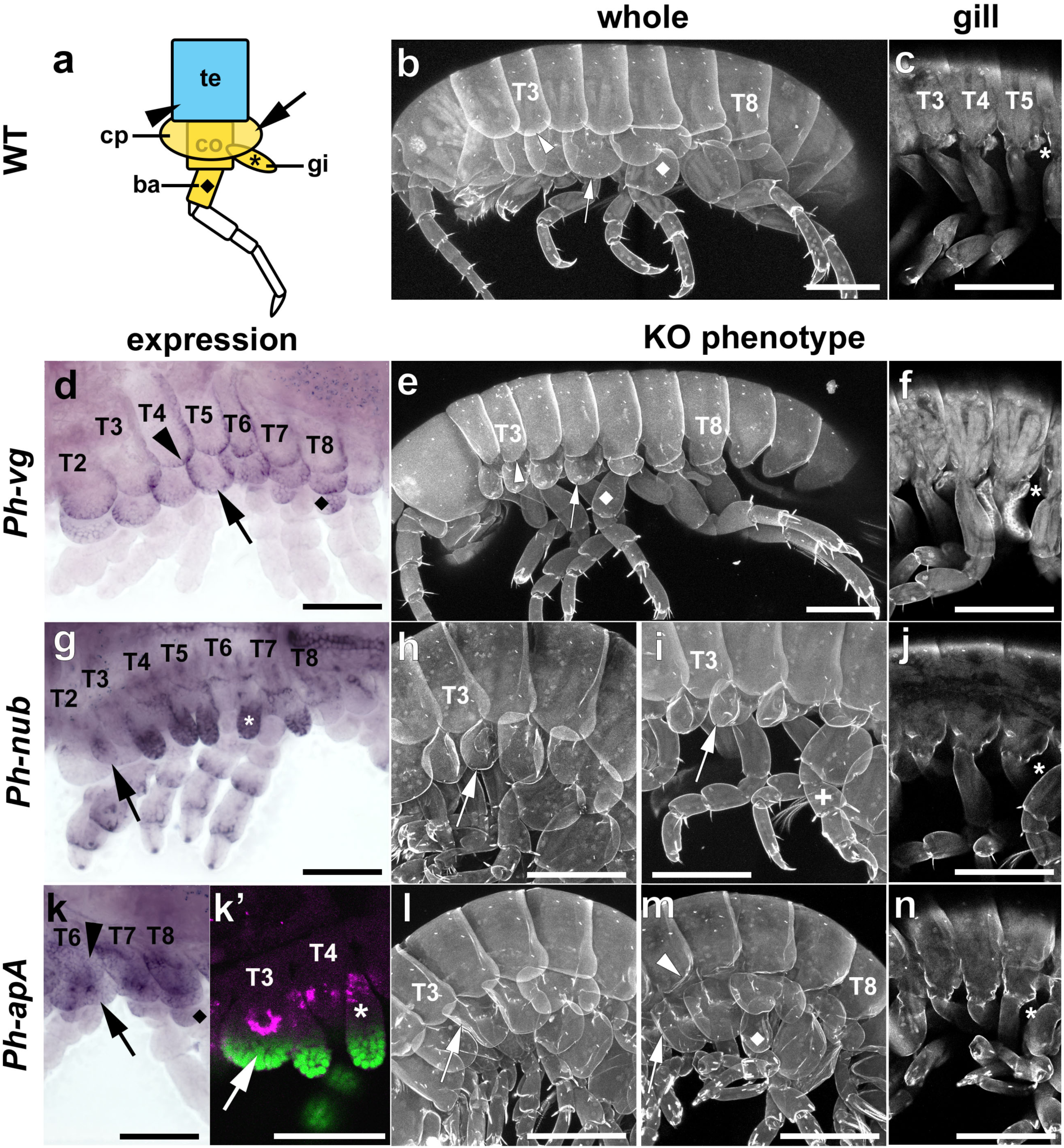
Expression and functional analyses of *vg, nub* and *ap* in *Parhyale*. **a**, Schematic of the dorsal-ventral body organization of *Parhyale.* te=tergum (arrowhead), co=coxa, cp=coxal plate (arrow), gi=gill (*), ba=basis (**♦**). Blue and yellow correspond to tergal and proximal leg tissues, respectively. **b, c**, WT whole (**b**) and optical section (**c**). **d-n**, *in situ* hybridization and CRISPR/Cas9 KO analyses for *vg* (**d-f**), *nub* (**g-j**), and *ap* (**k-n**). **k’**, optical section at the attachment of gill and coxal plate to coxa. *Ph-apA*: purple, DAPI: green. **+** in **i** indicates stunted leg. Scale bars 200*μ*m, except in **d**, **g**, **k** and **k’** 100*μ*m.

We first investigated the possible wing homologs in *Parhyale* via expression and functional analysis of *vestigial* (*vg*). The *vg* gene in insects is considered a critical wing gene because of its unique function in the ectoderm to orchestrate wing development^14,15^ and its potential to induce ectopic wings when overexpressed in certain contexts^16,17^. We and others have previously demonstrated that *vg* is quite powerful at identifying tissues serially homologous to wings in the wingless segments of insects^3,4,7,10,18,19^. In *Parhyale*, we found that the *vg* ortholog (*Ph-vg*, Extended Data Fig. 1) is expressed in the edge of the terga, as well as in parts of the proximal leg, including the edge of the coxal plate (cp) and part of the basis (Fig. 1d, SI movie 1). In addition, knocking out (KO) *Ph-vg* via CRISPR/Cas9 genome editing (Extended Data Fig. 2) resulted in deletion of the tergal edge, the entire cp, and the expansion of the basis (Fig. 1b and e, SI movie 2 and 3), validating the functionality of *Ph-vg* in these tissues. The penetrance of these *Ph-vg* KO phenotypes was very high, with the majority of G0 somatic cells displaying all relevant phenotypes (Extended Data Table 1 and 2). Intriguingly, although the gills of crustaceans have been previously proposed as crustacean wing homologs^20^, the gills of *Ph-vg* KO individuals remain intact (Fig. 1c and f, SI movie 2 and 3), suggesting that the gill of *Parhyale* may not be related to insect wings (however, the homology among crustacean gills requires further evaluation). Together, these findings suggest that both the terga (a dorsal tissue) and the proximal leg segments (homologous to insect pleural plates^21–23^ (Bruce et al. accompanying ms)) are the possible wing homologs in this crustacean.

**Figure 2.**
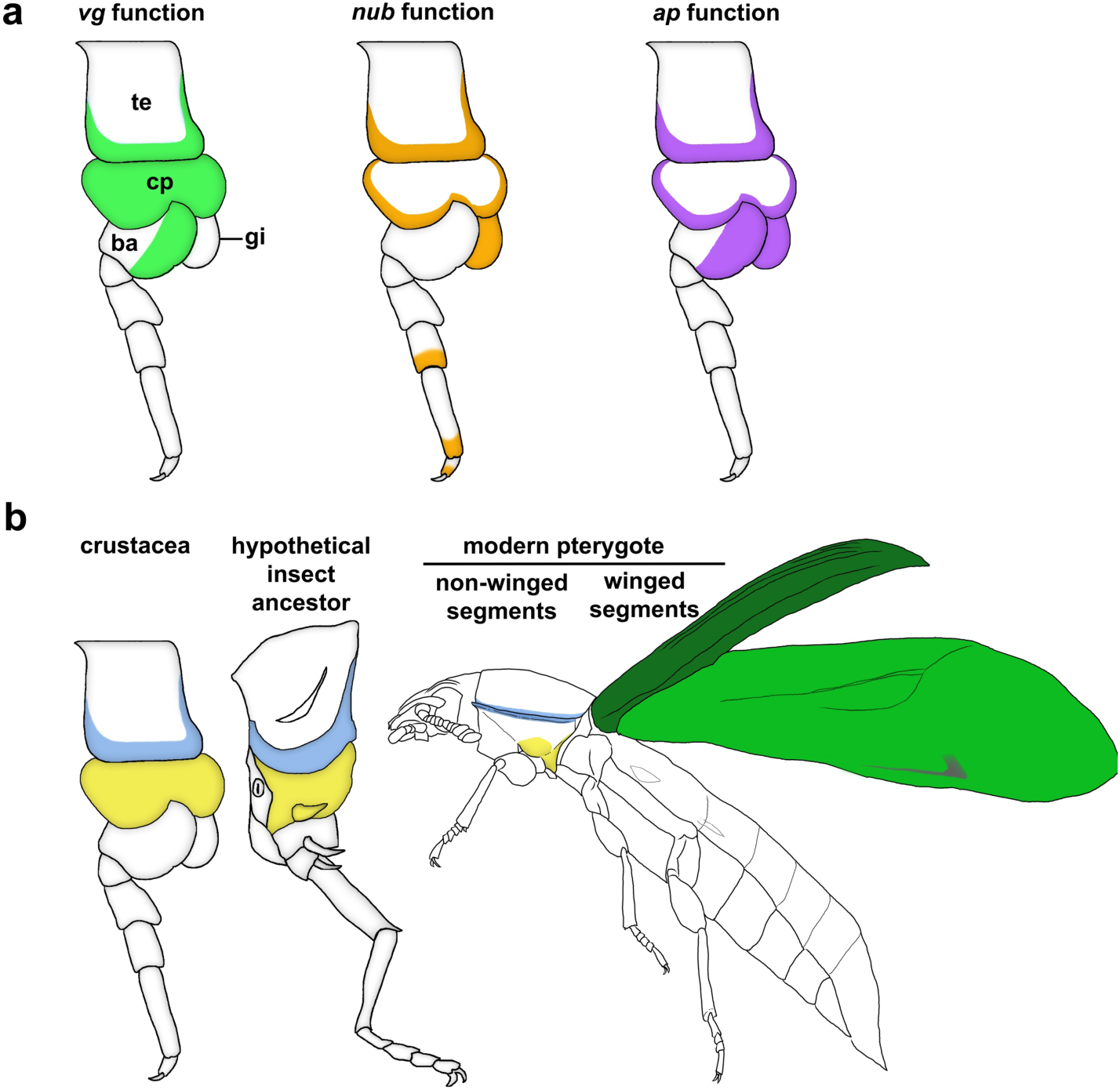
The proposed wing homologs of *Parhyale* and the evolutionary relationship among wing homologs. **a**, the functional domains of the three genes examined in this study. te=tergum, cp=coxal plate, gi=gill, ba=basis. **b**, the evolutionary relationship among wing homologs. The tergal edge (blue) and proximal leg segment (cp, yellow) are two possible crustacean wing homologs, in which two similar wing-like GRNs operate (left in **b**). These crustacean tissues correspond to the two proposed wing origin tissues; the crustacean tergum corresponds to the insect tergum, and the crustacean proximal leg segments to the insect pleural plates (middle in **b**). A similar situation can be found in the wingless segments of pterygote insects (right in **b**, tergal and pleural tissues are colored in blue and yellow, respectively). A wing-like GRN operates in both of these tissues, and they merge to form a complete wing upon homeotic transformation^4–6^, indicating that both tissues are wing serial homologs in the wingless segments of pterygotes (right in b). Through these observations, we propose that, prior to the evolution of insect wings, the apterygote ancestor of hexapods possessed two distinct tissues (of tergal and pleural nature), both of which had similar wing-like GRNs operating in them (middle in **b**). The evolution of pleural plates from the most proximal leg segment (*i.e.* subcoxa) has juxtaposed these two distinct tissues that rely on a similar GRN, which might have resulted in one functional unit of tissues (green) with a “cross-wired” GRN (ancestral wing GRN). In contrast, in the wingless segments, Hox genes have evolved to prevent this merger, maintaining the tissues serially homologous to wings as two separate sets (one of tergal and the other of pleural nature).

To further investigate the genetic overlap between the two *vg*-dependent tissues in *Parhyale* and insect wings, we analyzed two additional genes in *Parhyale, nubbin* (*Ph-nub*) and *apterous* (*Ph-ap*) (Extended Data Fig. 1). In insects, *nub* is strongly expressed in future wing tissues and loss of its function disrupts the development of wing-related tissues^5,24–26^. *ap* is expressed in the dorsal compartment of the wing as well as in the tergum in *Drosophila*^27–29^ and other insects^26,30^. In the *Drosophila* wing disc, *ap* acts as the dorsal selector to establish the DV organizer^27^. Both *nub* and *ap* have been used previously to identify wing homologs in crustaceans, leading to the identification of the gill as a potential wing homolog^20^. In *Parhyale, Ph-nub* is expressed strongly in the gill and weakly but broadly in the cp (Fig. 1g, SI movie 4). *nub* is also expressed in a ring pattern in each segment of the leg, which appears to match the *nub* leg segmental expression of various arthropods^31,32^ (Fig. 1g, SI movie 4). Upon knock-out, *Ph-nub* mutants consistently show loss of gills (Fig. 1j, Extended Data Fig. 3, Extended Data Table 1 and 2, SI movie 5). *Ph-nub* KO also causes reduction or curling of the cp, as well as leg miniaturization (Fig. 1h and i, SI movie 5). Interestingly, although the presence of *Ph-nub* expression in terga is somewhat ambiguous, we detected occasional mild tergal defects in *Ph-nub* KO individuals (Extended Data Table 2, Extended Data Fig. 4). Therefore, in *Parhyale,* the *nub* gene is essential for the proper formation of terga, as well as various leg components, including the gill, cp, and distal leg segments.

We previously identified the two classes of *ap* genes in arthropods, termed *apA* and *apB*, with *apA* being a dominant paralog during wing development^26,33^. The *ap* gene that has previously been tested in another crustacean appears to belong to the *apB* class^20,26^ (Extended Data Fig. 1). We identified two *ap* genes in *Parhyale,* corresponding to the two classes (*Ph-apA* and *Ph-apB*) (Extended Data Fig. 1). Our expression analysis revealed that *Ph-apA* is the relevant *ap* paralog for wing homolog identification, as *Ph-apB* is only expressed in the brain of *Parhyale* (Extended Data Fig. 5). In contrast, *Ph-apA* is expressed diffusely throughout the terga, cp, and basis (Fig. 1k, SI movie 6) and acutely where the cp and gill join the coxa (Fig. 1k’). *apA* KO (Extended Data Fig. 2 and 3, Extended Data Table 1) causes curling in the edge of the terga, the cp, and in the expansion of the basis (Fig 1l and m, SI movie 7). In addition, *apA* KO individuals are often missing the entire gill, even though *Ph-apA* expression is limited to the base of the gill (Fig. 1n, Extended Data Table 2, SI movie 7). Full deletion or severe reduction of all relevant tissues (the tergum, cp, and gill) was also observed with *apA* KO, but we were only able to recover one such individual (Extended Data Fig. 4). This low penetrance of severe KO phenotype might be due to high lethality when the majority of the somatic cells are *apA* KO. Taken together, our expression and functional analyses for the three “wing” genes in *Parhyale* have revealed that, although the expression and functional domains for each of these genes do not overlap completely, all three genes are critical for the formation of both terga and components of the proximal leg (Fig. 2a). Therefore, both of these two tissues are equally likely to be the wing homologs of crustaceans.

The debate on the origin of insect wings is like a pendulum that has been swinging back and forth between the two possible origin tissues for more than 200 years^3^. Previous molecular evidence of a wing GRN operating in crustacean gills^20^ strongly swayed this pendulum in the direction of a proximal leg origin of insect wing. Our identification of the wing-like GRN operating in both the terga and the proximal leg segments prior to the evolution of insect wings returns the swinging pendulum back to a neutral position, where either origin tissue can be implicated. As mentioned, there is a third direction for the pendulum to swing, namely toward a dual origin of insect wings^3^. Although not new, this idea has only recently been gaining momentum^4–7,34,35^. Below, we argue that the data presented here provide critical support for the dual origin model, and in combination with previous observations of wing serial homologs in wingless segments of insects, push the pendulum further in this new direction. First, the functional dependency of the two tissues on the wing-like GRN is not due to a common cell lineage of these tissues, as cell lineage tracing in the developing *Parhyale* embryo has identified that the tergum and the leg (including the most proximal components such as coxa) have different identities even early in development^36^. These data establish that the separation of these two lineages of tissues is deep in evolutionary time. Second, the presence of two separate sets of tissues per segment in *Parhyale*, both of which rely on a similar wing-like GRN, is reminiscent of the situation observed in the wingless segments of some insects where wing serial homologs are maintained as two separate tissues of tergal and pleural (i.e. ancestral proximal leg) nature^4–7,19^ (Fig 2). This similarity provides crucial support for the idea that the wingless segments of insects indeed reflect a plesiomorpic (ancestral) state for wing serial homologs, while the *bona fide* insect wing represents an apomorphic (derived) state that depends upon the contribution of two distinct tissues (*i.e.* a dual origin)^19^.

It is intriguing to speculate how a similar GRN has come to operate in the two distinct tissues (the terga and the proximal leg segments). Co-option of the GRN from one tissue to the other is a strong possibility that has recently been proposed^37^. Another possibility is shared ancestry between the terga and the proximal leg segments, as suggested in the accompanying paper (Bruce et al. accompanying ms). In either case, our data indicate that the wing-like GRN was already operating in both the terga and the proximal leg segments in the common ancestor of hexapods and crustaceans prior to the evolution of *bona fide* wings. It is also worth mentioning that the gill in *Parhyale* might not be homologous to insect wings, despite some genetic overlaps observed between these two structures. A previous study has demonstrated that the gill GRN has a larger overlap with the insect “respiratory GRN”^38^. Furthermore, here we showed that the gill lacks dependency on *Ph-vg*, a critical wing gene in insects. Therefore, the crustacean gill might be more homologous to the insect respiratory system than to the wing.

The evolutionary route from the two sets of wing homologs to the *bona fide* insect wing is still a mystery for future studies. The accompanying paper highlights one of the critical steps in the evolution of insect wings, *i.e.* the evolution of pleural plates (Bruce et al. accompanying ms). Bruce et al., provides compelling evidence supporting the idea that the most proximal part of the *Parhyale* leg (coxa) is equivalent to the insect subcoxa, a structure that has evolved into pleural plates in modern insects^21–23^. Considering our finding that similar wing-like GRNs are operating in both the terga and the proximal leg, the merger of the proximal leg into the body wall to form the pleural plates of insects could have been a key step in bringing these highly similar developmental modules closer together. Subsequently, this event may have caused a “cross-wiring” of the two similar GRNs operating in these two tissues, resulting in one fused tissue now functionally dependent on the merged GRN (*i.e.* an ancestral wing GRN) (Fig. 2b).

Although the evidence for a dual origin of insect wings is mounting^3–7,34,35^, this third hypothesis requires rigorous further testing from various fields. Recently established genetic techniques (such as the CRISPR/Cas9 genome editing used in this study) will allow us to delve deeper into the molecular basis underlying the evolution of insect wings. Meanwhile, the pendulum of the wing origin debate continues to attract more researchers to the unveiling of the origin and history of this evolutionarily monumental structure.

## Acknowledgements

We thank Nipam Patel and Heather Bruce for technical assistance and sharing their preprint, and Xavier Franch-Marro for the *Ph-vvl* clone. We also thank the Center for Bioinformatics and Functional Genomics (CBFG) and the Center for Advanced Microscopy and Imaging (CAMI) at Miami University for technical support, Shuxia Yi for technical assistance, and Takahiro Ohde, Ana Fernándes, David Linz, Nipam Patel, Heather Bruce, Arnaud Martin, and other members of the Tomoyasu and Patel labs for helpful discussion. This work is supported by the Miami University Faculty Research Grants Program (CFR) (to YT), the National Science Foundation (NSF) (IOS1557936 to YT), an NSF Graduate Research Fellowship (to CC-H), and EDEN: Evo-Devo- Eco Network (NSF-IOS0955517 to CC-H).

## Author Contributions

C. CH. and Y.T. conceived the experiments. C. CH. performed the experiments. C. CH and Y.T. analysed the data and wrote the manuscript.

